# Using neural networks to bridge scales in cancer: Mapping signaling pathways to phenotypes

**DOI:** 10.1101/324038

**Authors:** Eunjung Kim, Philip Gerlee, Alexander R.A. Anderson

## Abstract

Cancer is an evolving system subject to mutation and selection. Selection is driven by the microenvironment that the cancer cells are growing in and acts on the cell phenotype, which is in turn modulated by intracellular signaling pathways regulated by the cell genotype. Integrating all of these processes requires bridging different biological scales. We present a mathematical model that uses a neural network as a means to connecting these scales. In particular, we consider the mapping from intracellular pathway activity to phenotype under different microenvironmental conditions.

## I. BRIDGING SCALES IN CANCER: FROM PATHWAY TO PHENOTYPE

Much effort has been made to define molecular characteristics of cancer progression. This has led to the development of targeted therapies that successfully control cancer initially. The success, however, is generally short lived as drug resistance eventually emerges (1), primarily due to the multiscale aspects of cancer (2) and heterogeneity (3). Understanding how the resistance develops and can be delayed is one of major focus of cancer research (4). A strategy to deal with this resistance is to consider combination therapies with existing drugs. To develop more effective strategies, it is required to understand how selection by drugs affects intracellular pathway signaling and how the altered signaling affects the cell phenotype under dynamic and heterogeneous microenvironments.

Understanding the relationship between different intracellular molecules and relating their interactions to a cell phenotype is an overwhelm task considering the vast number of simultaneous interactions occurring within a cancer cell. Mathematical modeling may be a way forward with its ability to integrate these dynamics and to describe both linear and non-linear feedback between interacting molecules (5). A number of previous studies have used mathematical models for describing intra-cellular dynamics, utilizing a spectrum of techniques including Boolean, logic, artificial neural networks, and ordinary differential equations (6–26). Here, we focus on a neural network modeling approach as a means to connecting intracellular pathway and phenotype scale under different microenvironmental conditions.

## II. NEURAL NETWORK MODEL IN CANCER CELL SIGNALING

We utilized a neural network modeling approach to determine cancer cell phenotypes affected by both the microenvironment and intracellular pathway activity (Fig. 1). Feeding the microenvironmental variables to the hidden intracellular pathway layer determines the output of the network, cell phenotype (Fig. 1). In order to capture the fact that proteins in signaling pathway regulate other proteins, we allow for regulatory feedback between proteins. This modeling approach and preliminary results have already been published (12).

We model the mitogen-activated protein kinase (MAPK) pathway and the PI3K/AKT pathway (Fig. 1) since the pathways are known to mainly regulate cancer cell growth and death (27). Using the model, we first study normal signaling responses in different microenvironments, various concentrations of growth factors and death promoting signal. The normal cell network produces a profile of the proteins in Figure 2 in four different microenvironmental conditions, (i) low growth but high death, (ii) high growth and high death, (iii) high growth and low death, and (iv) low growth and low death. The predicted pro-growth outputs reflect different contexts. For example, the value is low in the micrioenvironment (i) and increases in growth favoring microenvironments (e.g., (iii)). The microenvironment (iv) pushes cells to an inactive state by decreasing all protein levels (Fig. 2).

**Figure 1.**
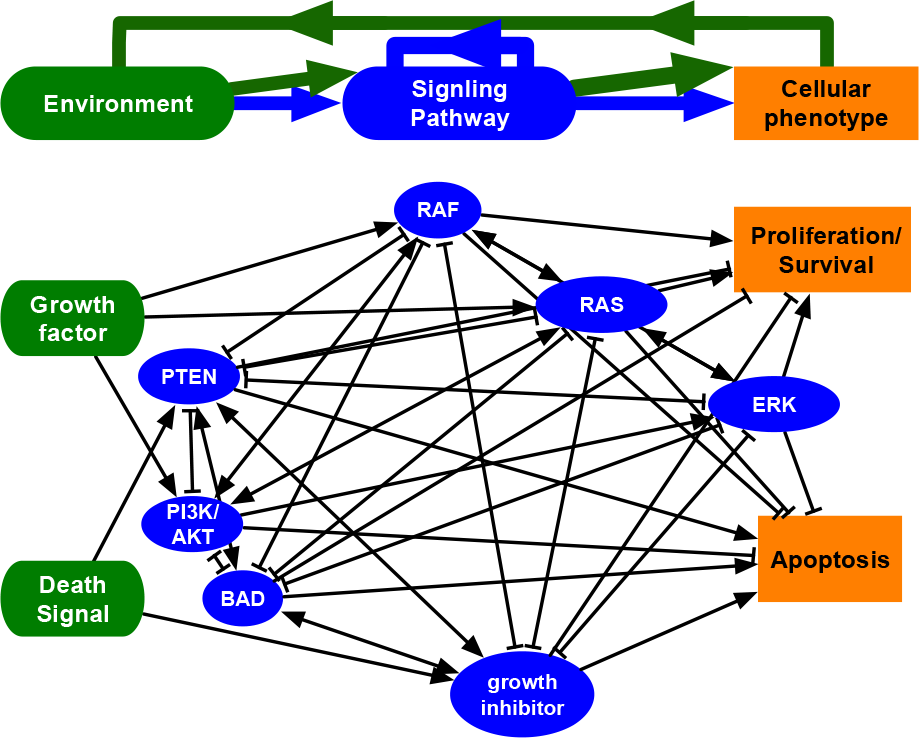
Artificial neural network. Microenvironmental variables (green) are in the input-layer and fed through the pathway (blue) and phenotype (orange), output layer. Connections between layers represent mapping from microenvironments to pathway and then pathway to phenotype.

**Figure 2.**
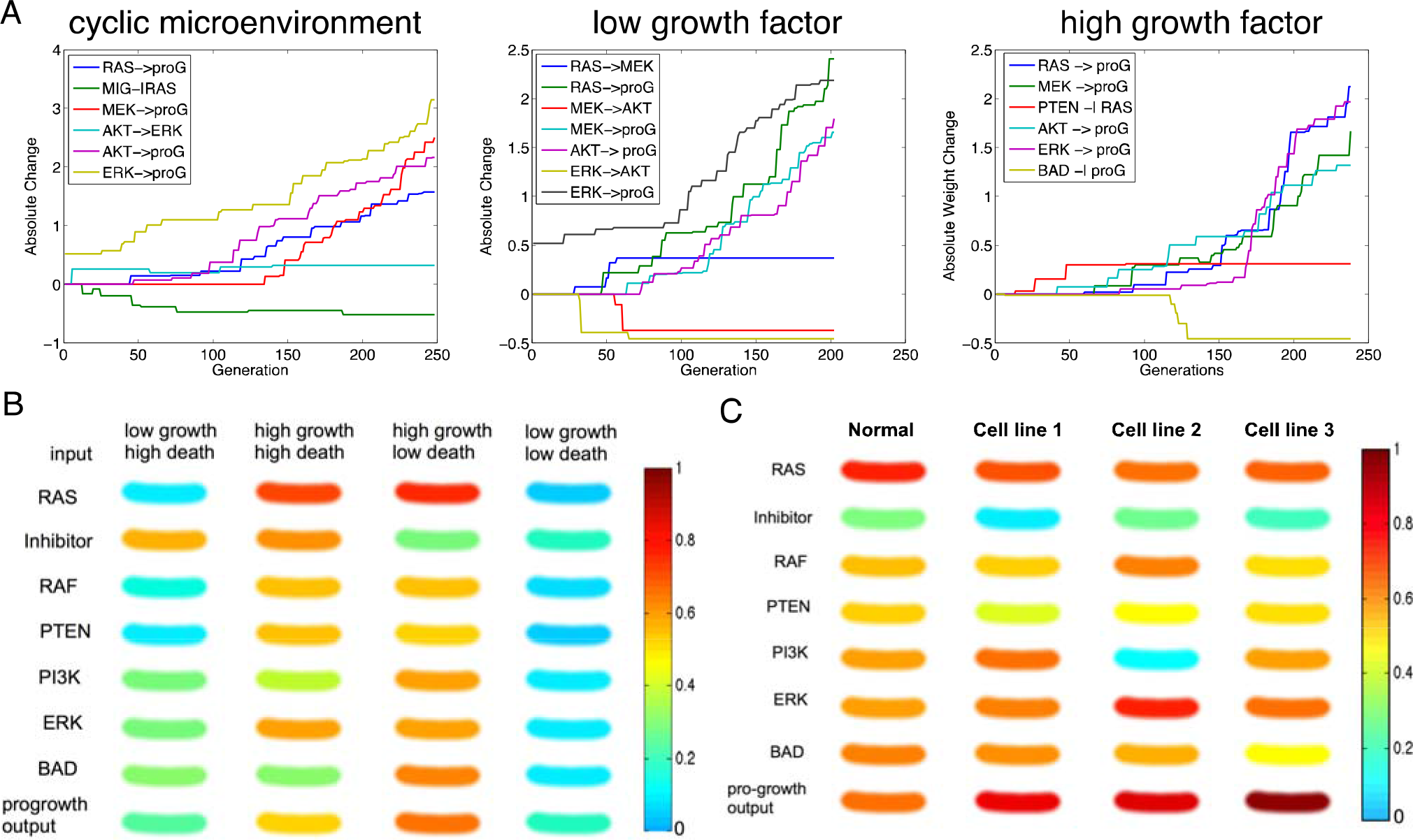
A: Changes of Weights from internal proteins to pro-growth output (proG) in a representative in silico cancer cell from cyclic, low, high growth factor microenvironment over 200 generations. B: Predicted protein activity and pro-growth output of normal cell in four different microenvironments indicated by the label on the top. C: Predicted protein activity as well as pro-growth output of normal and cancerous cell line (1-3) in microenvironment condition (iii), the high growth factor and low death factor condition.

Next, we evolve the normal network under different microenvironmental constraints to derive a cancer-signaling network. The network is evolved in three different microenvironments ((i) cyclic, (ii) low growth/high death, (iii) high growth/low death) to generate three different *in silico* cancer cell lines (cell line 1- cell line 3). We compare evolved weights of a representative *in silico* cancer cell from each microenvironment (Figure 2). The weights between internal proteins and pro-growth output seem to be increasing. We then compare the typical protein level of the normal network and three evolved cell lines (Fig. 2) in the high growth factor & low-death factor microenvironment (iii). Cell line 1 has a significantly lower than normal expression of the inhibitor protein (cyan) which suggests that it may harbor an inactivating mutation. Cell line 2 may have activating mutations in RAF and ERK as well as an inactivating mutation in PI3K. The expression of cell line 3 is only slightly different from that of a normal cell. Since cell line 3 was evolved in the grow-promoting environment, only a slight change in the network was sufficient to satisfy the given criteria (i.e., high pro-growth output).

## III. QUICK GUIDE TO THE METHODS

The network consists of two microenvironmental input nodes (growth factor and death signal), and two phenotype output nodes (pro-growth and pro-death), and several nodes that represent intra-cellular proteins. Only the internal nodes contain recurrent interactions. The rates of change of both protein level and phenotypes are determined as an additive linear combination of its neighbors weighted by interaction strength (**W**), which is described by

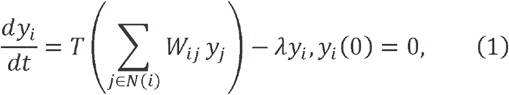

where *λ* is a decay rate. The transfer function *T* was considered 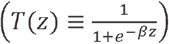 to account for saturation effect. Of note, *y*_1_ and *y*_2_ are external pro-growth and pro-death inputs.

To obtain a weight matrix, we use Monte Carlo simulation. A weight matrix is initialized with random numbers. With the initialized weight, we solve equation (1). Then we randomly select a weight element and perturb it. With this perturbed weight, equation (1) is solved again. The steady state solution with the perturbed weight and the one with the unperturbed weight are used to evaluate a pre-defined cost function. If the cost is closer smaller, the perturbation is accepted. Otherwise, the perturbation is discarded. The process is iterated until convergence.

### A. Derivation of normal cell weight matrix

In a normal cell, we assumed that the phenotype of the cell is directly regulated by microenvironmental cues. In other words, for a normal cell in a growth promoting microenvironment, it is more likely to reproduce. In a growth inhibiting or death promoting environment, a cell is less likely to divide and will have a higher chance of cell death. We model these phenomena using a cost function *C* = |*y*_1_ - *y*_*g*_| + |*y*_2_ - *y*_*d*_|, where *y*_1_ and *y*_2_ are given progrowth and pro-death inputs, respectively and *y*_*g*_ and *y*_*d*_ are pro-growth and pro-death outputs, respectively.

### B. Derivation of cancer cell weight matrix

A mutation is modeled by randomly perturbing each element of the normal cell weight matrix. To model higher growth rate of a cancer cell, a new cost function *C* = 1 − *y*_*d*_ was employed. Cell line 1 is evolved in condition (i), cell line 2 in (ii), and cell line 3 is evolved in the condition (iii).

## IV. OTHER APPLICATIONS

Artificial neural networks have been used as a machine learning approach for the detection of heart abnormalities (28–30) and cancer prediction (31–41). Our approach was also previously used to investigate the impact of the microenvironment on cancer growth and evolution (42–46), where the neural networks were embedded into an individual cell based model, hybrid cellular automata model, which allows for mutation in the cancer cells and subsequent selection by the microenvironment. A recent study utilizing similar approach highlighted the impact of targeted therapy on cell signaling heterogeneity (47).

